# Developmental and physiological impacts of pathogenic human huntingtin protein in the nervous system

**DOI:** 10.1101/2024.08.30.610525

**Authors:** Tadros A. Hana, Veronika G. Mousa, Alice Lin, Rawan N. Haj-Hussein, Andrew H. Michael, Madona N. Aziz, Sevinch U. Kamaridinova, Sabita Basnet, Kiel G. Ormerod

**Affiliations:** Middle Tennessee State University, Biology Department, Murfreesboro, Tennessee, United States of America; Brown University, Neuroscience Graduate Program, Warren Alpert Medical School, Providence, Rhode Island, U.S.A.

**Author notes:** Correspondence and requests for materials should be addressed to K.G.O.

**Keywords:** Drosophila, Huntington’s Disease, organelle trafficking, neuromuscular junction, aggregation

## Abstract

Huntington’s Disease (HD) is a neurodegenerative disorder, part of the nine identified inherited polyglutamine (polyQ) diseases. Most commonly, HD pathophysiology manifests in middle-aged adults with symptoms including progressive loss of motor control, cognitive decline, and psychiatric disturbances. Associated with the pathophysiology of HD is the formation of insoluble fragments of the huntingtin protein (htt) that tend to aggregate in the nucleus and cytoplasm of neurons. To track both the intracellular progression of the aggregation phenotype as well as the physiological deficits associated with mutant htt, two constructs of human HTT were expressed with varying polyQ lengths, non-pathogenic-htt (Q15, NP-htt) and pathogenic-htt (Q138, P-htt), with an N-terminal RFP tag for *in vivo* visualization. P-htt aggregates accumulate in the ventral nerve cord cell bodies as early as 24 hours post hatching and significant aggregates form in the segmental nerve branches at 48 hours post hatching. Organelle trafficking up-and downstream of aggregates formed in motor neurons showed severe deficits in trafficking dynamics. To explore putative downstream deficits of htt aggregation, ultrastructural changes of presynaptic motor neurons and muscles were assessed, but no significant effects were observed. However, the force and kinetics of muscle contractions were severely affected in P-htt animals, reminiscent of human chorea. Reduced muscle force production translated to altered locomotory behavior. A novel HD aggregation model was established to track htt aggregation throughout adulthood in the wing, showing similar aggregation patterns with larvae. Expressing P-htt in the adult nervous system resulted in significantly reduced lifespan, which could be partially rescued by feeding flies the mTOR inhibitor rapamycin. These findings advance our understanding of htt aggregate progression as well the downstream physiological impacts on the nervous system and peripheral tissues.

## Introduction

Huntington’s Disease (HD) is an inherited, autosomal dominant neurodegenerative disorder affecting 1/10,000-1/20,000 people in the US annually ^1^. HD is part of a broader family of polyQ disorders, which include Machado-Joseph Disease, the five spinocereberal ataxis types (1, 2, 6, 7, 17), spinal-bulbar muscle atrophy X-linked type 1 and dentatorubral pallidoluysian atrophy ^2,3^. PolyQ disorders are linked to a pivotal expansion of a trinucleotide cytosine–adenine–guanine (CAG) repeat encoding a polyQ (Q: amino acid glutamine) tract in the coding region of causative genes ^4^. Clinically, polyQ disease patients experience progressive increases in loss of motor control, cognitive decline, and psychiatric disturbances ^5,6,7^ In each disease-case, expansion of the polyQ region within the protein leads to protein aggregation; and although many of the disorders develop from mutations of unrelated genes, they share molecular and clinical characteristics ^8,9,10,11,12^.

HD is linked to a CAG expansion within exon 1 of the IT15/Huntington gene ^13^. Expansion of the CAG repeat leads to expansion and increased length of the PolyQ region of the huntingtin protein (htt); greater expansion leads to earlier disease onset and greater penetrance of disease pathology ^14,15,16^. While the htt protein is ubiquitously expressed, expanded CAG homorepeat mutations exert their pathology almost exclusively on neuronal tissue ^17^. Critically, the pathophysiology of HD is linked to protein fragmentation leading to the formation of insoluble inclusion bodies (aggregates). These aggregates are observed throughout neurons commonly in the nucleus, cytoplasm, and axons ^18,19^. In its early stages, HD displays preferential expression in the striatum, specifically increased neurodegeneration targeting medium spiny neurons in stages designated 1 through 4 ^20–22^. Interestingly, only 1-4% of striatal neurons display aggregation, potentially due to the neuroprotective effects of the full length htt protein ^23^. In contrast, aggregation is observed more often in other neuron types such as cortical and cerebellar when compared to striatal ^24^. Importantly, N-terminal htt fragments have been found to aggregate and exert pleotropic toxic effects on neurons ^18,25–29^. In particular, proteolysis of htt at the caspase-6 cleavage site is a critical event in mediating neuronal disfunction and neurodegeneration ^30^. Of the numerous other defects, htt aggregation is well known to lead to deficits in axonal trafficking ^31,32^. However, understanding the downstream implications of htt aggregation in neuronal structure and neuromuscular structure and function remains poorly understood. Therefore, understanding the developmental proliferation of htt fragment aggregation and putative downstream implications will lead to a better comprehension of HD progression and pathophysiology.

Herein, we exploited the extensive genetic and molecular toolkits in *Drosophila* to investigate the progression of htt fragment aggregation using pathogenic and non-pathogenic polyQ constructs truncated at the caspase-6 cleavage site with an RFP tag for *in vivo* visualization to accurately model the behavior of the mutant htt. This model facilitates *in vivo* htt-aggregation tracking in the central and peripheral nervous systems to elucidate the progression of their formation. Combining fluorescently tagged htt-constructs and fluorescently tagged-organelles (synaptotagmin-1 and a Brain Derived Neurotrophic Factor-GFP generated here), demonstrated profound impacts of htt-aggregates on the trafficking dynamics of synaptic vesicles and dense core vesicles in vivo. Htt-aggregates did not appear to alter neuromuscular junction morphology, however, excitation-contraction coupling in larvae expressing the pathogenic version of htt proteins was severely impaired, which also manifested as locomotor deficits. To generate greater longitudinal studies, the htt-constructs were successfully expressed in the adult wing, which showed a similar aggregation profiles to larvae. Lastly, to explore the cellular pathways underlying the proliferation of htt-aggregates, the mTOR pathway was targeted using rapamycin in adults, resulting in a significant increase in lifespan of the flies expressing pathogenic htt in the nervous system.

## Materials and Methods

### Husbandry

*Drosophila melanogaster* were cultured on standard medium at 22°C at constant humidity on a 12:12 light:dark cycle. Fly lines were obtained from the Bloomington *Drosophila* Stock Center (BDSC), see Table 1. UAS-htt-Q15, UAS-htt-Q138, and UAS-Syt1-GFP fly lines were generously provided by Troy Littleton, Massachusetts Institute of Technology.

**Table 1:** Fly lines, genotypes, and abbreviations used throughout investigation.

| Full genotype | Abbreviation | BDSC Stock # | Reference |
| --- | --- | --- | --- |
| UAS- HttQ15-mRFP/+ | NP-htt | - | 32,33 |
| UAS- HttQ138-mRFP/+ | P-htt | - | 32,33 |
| Elav-Gal4/+ | Elav | 458 | 34 |
| Elav-Gal4/+ ; UAS- HttQ15-mRFP/+ | Elav > NP-htt | - | 32,33 |
| Elav-Gal4/+ ; UAS- HttQ138-mRFP/+ | Elav > P-htt | - | 32,33 |
| OK6-Gal4/+ | OK6 | 64199 | 35 |
| UAS-Syt1-GFP/+ | Syt1-GFP | - |  |
| UAS-BDNF-GFP/+ | BDNF-GFP | - |  |
| UAS-Mito-GFP | Mito-GFP | 95270 | 36 |
| Canton. S. | Canton. S. | 6365 | 37 |
| Motor Neuron Is-Gal4 | MN-Is | - | 38 |
| Motor Neuron Ib-Gal4 | MN-Ib | 40701 | 38 |
| appl-Gal4 | appl | 32040 | 39 |

### Generation of transgenic construct

cDNA of preproBDNF-eGFP (BDNF-eGFP) was subcloned into pUAST vector. cDNA of mouse preproBDNF was kindly provided by Prabhodh Abbineni (Loyola University Chicago). Microinjection of constructs was performed by BestGene Inc. to generate transgenic *Drosophila* strains.

### Larval htt aggregation imaging

#### Egg-laying assay

Three (3) adult males and three (3) adult females were isolated the day of eclosion and mated in a separate food vial. On day 3, the flies were transferred to an egg-laying assay which included a 35 mm petri dish with grape agar, and yeast paste. Flies were left for 24 hours to lay eggs, and afterwards, unhatched eggs were transferred onto a new food plate ^40^. Newly hatched larvae were collected and isolated on separate dishes at two-hour intervals. Larval age was measured as hours after hatching (HAH).

#### Transcutaneous (intravital) imaging (24, 48, and 72 HAH)

Larvae were transferred from food vials to microscope slides. Clear tape was applied to slides, creating a small channel with dimensions matching the height and width of the larvae. Larvae were then coated with a layer of glycerol and a cover slip was placed on top. Slides were refrigerated at 4°C for three-to-five minutes to fully immobilize the larvae prior to imaging.

#### Dissected preparations (96 and 120 HAH)

Early and late third-instar larvae were dissected in HL3.1 calcium-free saline with the following composition (in mM): NaCl: 70; KCl: 5; CaCl_2_:1.5; MgCl_2_: 4; NaHCO_3_: 10; Trehalose: 5; Sucrose: 115; HEPES: 5, (pH = 7.18) ^41^. All images were taken with a Nikon Eclipse microscope, equipped with a Lumencor Fluorescence Light Engine, Hammamatsu Orca-fusion camera, and processed using Nikon Elements software. All images were taken with a 60x water-immersion objective. ***Analysis:*** To determine aggregate thresholds, RFP-positive puncta were recorded live from the VNC of five (5) different Elav > NP-htt larvae at 25 frames per second (fps). From the five (5) animals, a total of 25 000 µm of nerves were assessed to identify the largest motile aggregate, found to be 1.8 µm^2^. Consequently, for analysis the size threshold was set at 150% of 1.8 µm^2^ (2.7 µm^2^) and the fluorescence intensity threshold to 2 SD above background. ***First-and second-instar image analysis.*** All data were obtained using the ROI feature on Nikon Elements. The ventral nerve cord (VNC) was identified by the presence of the neuropil. Fluorescence intensity was extrapolated by manually tracing around the VNC. For segmental nerve branch (SNB) analysis, at least 500 µm of nerves were imaged from each preparation, and aggregates forming in the motor neuron (MN) axons were manually selected and traced. Number, average fluorescence intensity, and size were all measured from the aggregates in each ROI and measurements were exported to Excel. ***Early-and late-third instar (96 and 120 HAH).*** All data were obtained using the ROI feature on Nikon Elements. The VNC was identified by the presence of the neuropil or by tracing the SNBs to their origin point. SNBs were imaged 450 µm downstream of the brain, and each preparation had a minimum of ten (10) individual nerve branches to be analyzed. Number, average fluorescence intensity, and size were all measured from the aggregates in each ROI and measurements were exported to Excel.

#### Morphological Measurements

Twenty (20) randomly selected wandering third-instar larvae from each genotype were selected and rinsed in deionized water prior to being measured. Larvae were pinned on the posterior and anterior ends using minuten pins and stretched until abdominal contractions were no longer observed. Length, at the longest points, and width, at the widest points, were recorded using a dissection microscope equipped with an objective reticule. Area was calculated as the product of length times width.

#### Organelle trafficking

Wandering larvae were isolated from their respective vials and dissected in standard HL3.1 saline. Live imaging of 3^rd^ instar larval SNB was performed using UAS-Syt1-GFP (SV) and UAS-BDNF-GFP (DCV) in both the pathogenic-(P) and nonpathogenic-(NP) htt backgrounds. 90 second videos were recorded at 4 fps on a Zeiss Axio Imager 2 equipped with a spinning-disk confocal head (CSU-X1; Yokagawa) and ImagEM X2 EM-CCD camera (Hammamatsu). An Olympus LUMFL N 60X water-immersion objective with a 1.10 NA was used to acquire images. All videos were acquired at the same fps, and laser intensity. Velocity 3D Image Analysis software (PerkinElmer) was used to analyze images. Kymographs were extracted and analyzed using Kymograph Clear and Kymograph Direct.

#### Immunohistochemistry

Wandering third-instar larvae were dissected in hemolymph-like saline HL3.1. The larvae were fixed for three minutes in 4% paraformaldehyde and subsequently washed in phosphate-buffered saline (PBS), with 0.05% Triton X-100 (PBST), and blocked with 5% BSA. Larvae were then incubated with antibodies in PBST at room temperature for 2 hours and washed three times for 15 min in PBST. Finally, larvae were mounted in a medium containing DAPI (ab104139). Antibodies used for this study include the following: DyLight 649 conjugated anti-horseradish peroxidase (HRP), 1:500 (catalog #123–605-021, Jackson ImmunoResearch) and the filamentous actin probe, phalloidin conjugated to Alexa Fluor™ 488 (A12379, Thermo). Immunoreactive proteins were imaged on the Nikon fluorescence microscope (see above) using either 20x air or 60x-water immersion objectives and processed using Nikon Elements software.

#### Neuromuscular junction analysis

NMJ analysis was performed using a horseradish peroxidase (HRP) stain. Innervation length was obtained using a distance measurement feature on Nikon Elements to measure the extent of innervation on the surface of muscle fiber 4. Bouton number and fluorescence values for each genotype were calculated using regions of interest (ROI) tool in Nikon Elements, by manually selecting and tracing HRP-positive boutons. Average bouton fluorescence and area from each ROI were exported from the software and bouton number and area were subsequently exported to excel.

#### Muscle Examination

Length and width of individual muscles were quantified using the distance measurement feature on Nikon Elements. Muscles from abdominal segments A3-A5 were used. Muscle length was measured along the middle of muscle fiber 4, using the midpoint of the muscle insertion point on either end of the muscle. Muscle width was measured across the midline of the muscle, equidistant from both muscle insertion points. The product of both length and width were used to compute muscle area. Muscle sarcomere length was measured using the intensity profile function in Nikon Elements software to produce a sinusoidal graph of anti-actin fluorescence intensity. The length of the sarcomere was then measured as a single sinusoidal wavelength. Moreover, I-band was obtained by measuring the width of each peak from the first significant decrease in fluorescence ^40^. Fixed dissections were mounted onto microscope slides with DAPI mounting media (ab104139). Nuclei area was measured from overlayed images of DAPI and anti-actin using a region of interest (ROI) measuring tool on Nikon Elements. Subsequent fluorescence intensity, number, and area of each ROI was then exported to Microsoft Excel. Inter-nuclei distance was obtained by taking the three most central nuclei and measuring the three closest nuclei to them.

#### Muscle force recordings

Force recordings were conducted as described in Ormerod et al, 2022 ^42^. Briefly, third instar larvae were pinned dorsal side up and a lengthwise cut was made along the dorsal midline. Submerged in modified HL3.1 saline (4mM Mg and 1.5 mM Ca^2+^), the preparation was eviscerated, and the CNS was removed. Segmental nerve branches were stimulated using a suction electrode and A.M.P.I. Master 8 stimulator. Force recordings were acquired using the Aurora Scientific 403A Force Transducer System, which includes a force transducer head stage, amplifier, and digitizer. Contractions were elicited using 5x10^-4^ s impulses for 600 ms, with an interburst duration of 15 s, with varying intraburst stimulation frequencies (1-150 Hz). Every preparation was visually inspected to ensure at least 5 abdominal segments were stimulated. Digitized data was acquired using Aurora Scientific Software, Dynamic Muscle Acquisition (DMCv5.5). Digitized data were imported and processed in MATLAB using custom code ^42^.

#### Electrophysiology

Wandering third instar larvae were isolated from their vials and rinsed with DI H2O. Standard dissection protocol was followed ^43^, and preparations were bathed in HL3.1 (0.25 mM Ca^2+^) ^41^. Larvae were pinned dorsal side up and motor nerve branches were stimulated (A.M.P.I. Master 8) using a suction electrode following removal of the CNS. All animals were stimulated at 0.2 Hz and intracellular voltage was recorded using a sharp recording electrode (40-60 MΩ) from muscle fiber 6, in abdominal segments A3-A5. Voltage traces were digitized using a Minidigi 1B (Molecular Devices) and visualized using Axoscope (Molecular Devices) Software. Excitatory junctional potentials (EJPs) and miniature excitatory postsynaptic potentials (minis) were quantified using ClampFit, and data were explored to Microsoft Excel for data compilation, and subsequently to GraphPad prism for statistical analysis and figure making.

#### Larval Crawling

Third instar wandering larvae were isolated from fly vials and washed seven times in DI water. Groups of 10 larvae from the same genotype were transferred to the center of a 1% agar plate for a 5-minute crawling session. 10 separate recordings of 10 larvae per recording were taken for each genotype assessed. Recordings of 10 larvae per genotype were made using an infrared camera housed inside a 24”x24”x24” black box. Videos were processed using SpinView and subsequently converted to an Interoperatory Master Format (IMF) file. Caltech Multiple Fly Walking Tracker (CTRAX) software extracted a CSV file with X and Y coordinates and heading in radians. Excel scripts analyzed the data for distance per frame, displacement, velocity per 0.5 seconds, and angular velocity, revealing genotype-specific locomotor traits.

#### Adult Lifespan Assay

For each genotype, 400 total flies were tracked daily from eclosion to death. The 400 flies were divided into separate vials of 20 flies, each with 10 female and 10 males, isolated upon the day of eclosion. Once per week, the flies were transferred to fresh food vials. Each vial was incubated at 22°C and examined daily by counting the number of nonviable flies and the sex.

#### Rapamycin

Standard fly food was prepared with Rapamycin (Medchemexpress) dissolved in 1 mL of dimethylsulfoxide (DMSO). 400 adult P-Htt flies were separated into 20 separate vials containing 10 males and 10 females. Adults were isolated within a 24-hour window post eclosion and designated as day 1 old. Flies were transferred to rapamycin-containing food as indicated in the legend of Figure 10. Flies were tracked every day and transferred to fresh food vials every five days. The number and sex of nonviable flies each day were counted, and the data were input into Microsoft Excel.

#### Wing imaging

Imaging was performed every 5 days for 50 days. At each timepoint, adult flies were removed from food vials and transferred to a CO_2_-pad. Wings were removed via cutting the hinges and visualized under a dissection microscope. The wings were mounted on a microscope slide with glycerol. Wings were imaged using the same Nikon microscope as above, and a 60x oil objective. The entirety of L1 of each wing was imaged and then stitched together using Nikon Elements Software. After imaging, Nikon Elements Software was used to select non-motile aggregates using a threshold of 2.7 µm ^2^ and a fluorescence intensity 2 SD above the background.

### Statistical analysis

GraphPad Prism 10.3.0 was used for statistical analysis. Appropriate statistical metrics were performed for each dataset along with the F, and P statistics. Statistical comparisons were made with controls unless noted. Appropriate sample size was determined using a normality test. Data are presented as the mean ± SEM unless otherwise stated (*p<0.05, **p<0.01, ***p<0.001, n.s. = not significant).

## Results

### Effects of Htt-aggregation on development and gross morphology

Given that huntingtin (htt) intracellular protein aggregation is a progressively occurring neurodegenerative disease, the effects of pathogenic versus non-pathogenic human htt expression on the growth and development of *Drosophila melanogaster* were initially investigated. Using the UAS/Gal4 system, *Drosophila* expressing either a nonpathogenic polyQ tract of 15 repeats (NP-htt), or a pathogenic polyQ tract of 138 repeats (P-htt) corresponding to a juvenile form of HD, within the human Htt gene, under the control of UAS were investigated ^33^. Both constructs contain a fused N-terminal red fluorescent protein (RFP) to enable *in vivo* visualization and analysis of the formation and localization of htt. Given that proteolysis of full length htt leads to aggregate formation, we employed a truncated form of htt at the caspase-6 cleavage site comprising the initial 588 amino acids to accurately model the aggregation phenotype ^30,33^. To selectively examine htt progression within the nervous system (NS), both transgenic lines were crossed with the neuronal driver Elav-Gal4 (Elav). To fully characterize the effects of P-htt expression on the growth and development of *Drosophila* initially, each stage of development was examined. Initially, an egg-laying assay was conducted using 3 males and 3 females (Fig 1). After 24 hours, the total number of eggs were counted for both transgenic lines expressed in the NS (Elav > NP, Elav > P) as well as control lines (Elav, UAS-NP-htt, and UAS-P-htt); no statistical differences were observed (Fig 1 A, C. One-way ANOVA, F=1.3, P=0.31). Next, the number of eggs that hatched into first instars were counted; no statistical differences were observed between genotypes (Fig 1 D, One-way ANOVA, F=0.87, P=0.49). Subsequently, 20 third instar larvae for each genotype were collected and placed into separate vials (Fig 1B). No significant differences were observed in percentage of larvae that underwent pupation between the 5 genotypes (Fig 1 E, One-way ANOVA, F=3.41, P=0.08). Lastly, the percentage of larvae eclosed revealed significant differences between Elav > NP-htt and Elav > P-htt, as well as between Elav > NP-htt and NP-htt (Fig 1 F, One-way ANOVA, F=4.65, P=0.008). Taken together, expression of P-htt in the nervous had no impact on development until the adult stage.

**Fig 1.**
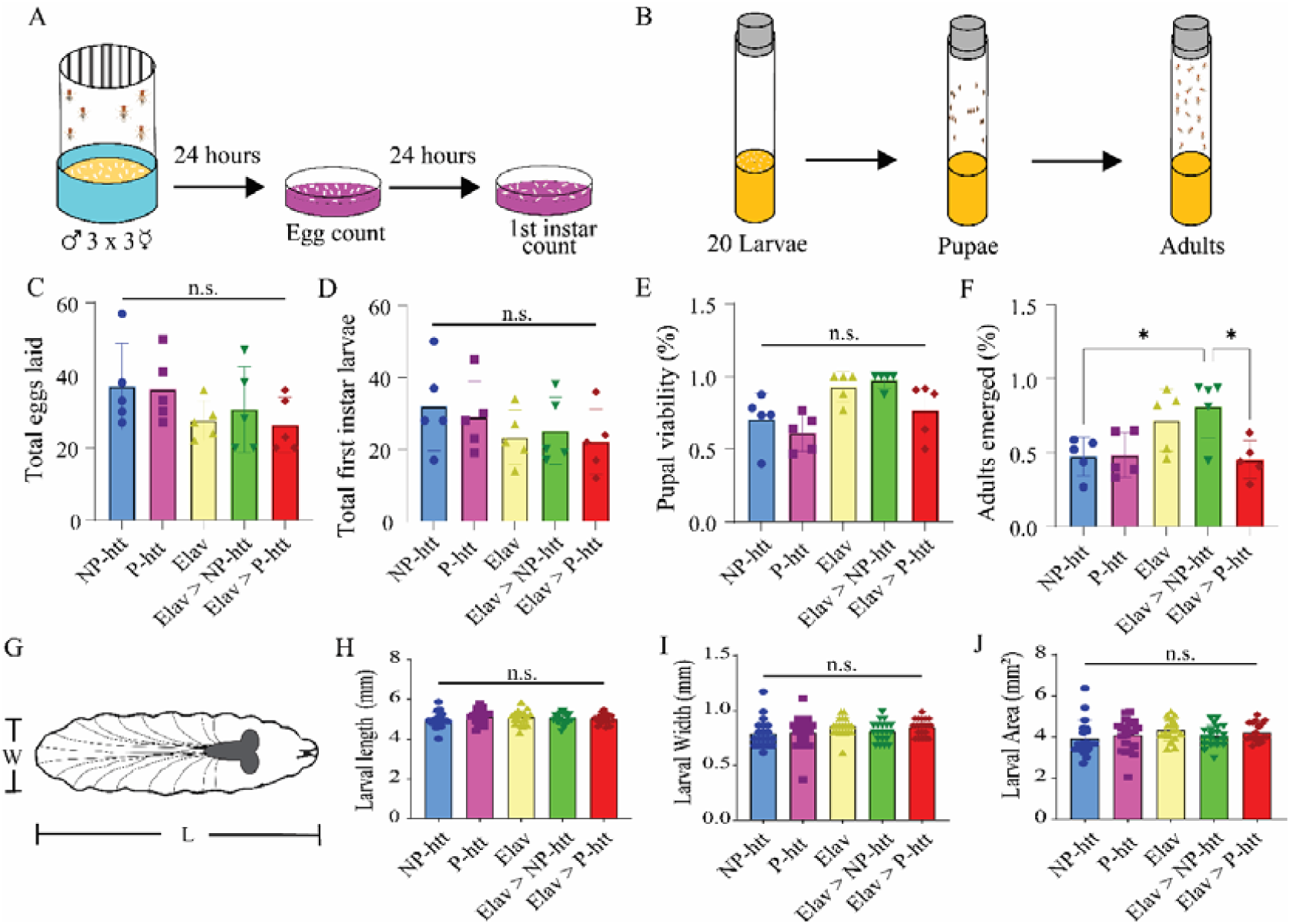
Pathogenic htt protein aggregation does not impact larval morphology or development. **A.** Diagram of egg laying assay. Eggs were collected from a cross between 3 males and 3 females after 24 hours and emerging first-instar larvae were collected 24 hours afterwards. **B.** Diagram of larval development assay. 20 larvae were transferred onto food vials and tracked until pupation and adulthood. No significant differences were observed from the 5 different genotypes investigated from the total eggs laid **(C),** total first instar larvae counted **(D)** or pupal viability **(E). F.** Percentage of adults that emerged from each crossed showed significant differences. **G.** A schematic depicting a third-instar larva being measured for length and width. No significant differences were found in larval length **(H)**, width **(I)**, or area **(J)** across the 5 genotypes investigated. * indicates P>0.05 from a One-way ANOVA.

To further assess the potential impacts of htt-aggregation on growth and development, an assessment of third instar larvae expressing P-htt or NP-htt, gross larval morphology was conducted (Fig 1 G). Twenty randomly selected third instar larvae from each of the 5 genotypes were measured and no significant differences in larval length (Fig 1 H, One-way ANOVA F=0.75, P=0.56), width (Fig 1 I, One-way ANOVA, F=1.33, P=0.26), or area (Fig 1 J, One-way ANOVA, F=1.13, P=0.35) were observed. These results suggest that htt-expression in the nervous system does not impact developmental timing or gross morphology.

### Larval Htt-aggregation assay

Having demonstrated that htt-expression in the NS does not impact developmental timing or gross morphology, an examination of cellular P-htt progression within individual neurons was conducted. Given the semi-transparent nature of first and second instar larvae, we were able to perform transcutaneous imaging from fully intact, undissected animals ^44^. To precisely track the progression and proliferation of htt-aggregation, eggs were collected every 2 hours and sorted into separate fly vials facilitating a precise 2-hour window for developmental tracking based on hours after hatching (HAH). First instar larvae were imaged transcutaneously for htt expression in the NS using the Elav driver, which expresses prolifically in motor neurons ^34^. In Elav > NP-htt larvae at 24 HAH, no quantifiable RFP expression was observed in the ventral nerve cord (VNC) when measuring the average fluorescence intensity of the entire VNC relative to background (VNC, Fig 2 Ai, C). At 48 HAH, a significant increase in RFP fluorescence was observed in Elav > NP-htt larvae, but no further significant increase was observed for 72, 96, or 120 HAH compared to the 48 H timepoint (Fig 2 Aii-v, C, One-way ANOVA, P>0.05). For Elav > P-htt larvae, quantifiable RFP fluorescence was observed in the VNC 24 HAH, significantly greater than Elav > NP-larvae of the same age (Fig 2 Avi, C, One-way ANOVA, F= 85.99, P<0.0001). A day later, in 48 HAH Elav > P-htt larvae, significantly more htt-aggregation was observed compared to 24 H (Fig 2 Avi-vii, C, P<0.01). At 72 HAH, significantly more fluorescence was observed compared to both 24 and 48 HAH (Fig 2 Avi-viii, C, P<0.01); however, 96 and 120 HAH timepoints did not show any differences compared to 72 HAH (Fig 2 Aviii-x, C). Thus, in Elav > P-htt larvae, htt-aggregation steadily accumulated in the VNC from 0-72 HAH, however, no further htt-proliferation was observed after 72 HAH (Fig 2 Avi-x, C). Conversely, quantifiable RFP fluorescence was not observable until 48 HAH for Elav > NP-htt larvae, and no significant further increases in fluorescence was observed thereafter (Fig 2 Ai-v, C).

**Fig 2.**
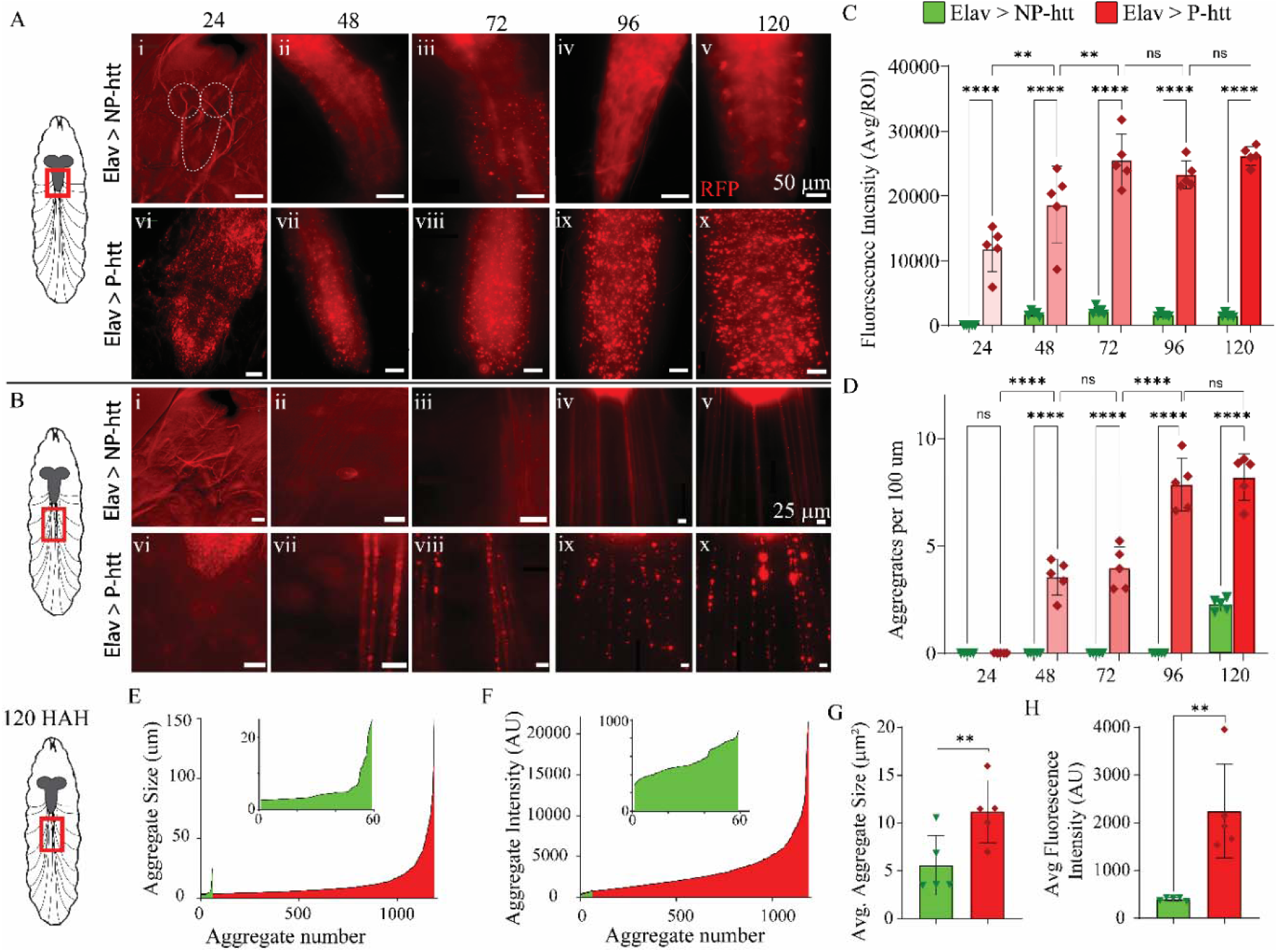
Expanding the PolyQ repeat region causes a progressive proliferation of htt aggregates throughout larval development. **A.** *Left:* Schematic depicting site of imaging on a third-instar larva. Intravital images (24, 48, 72 Hours after Hatching) and dissected preparations (96 and 120 Hours after Hatching) of Ventral Nerve Cords (VNC). First row: representative images of larvae expressing Elav>NP-*htt* from each time point. Second row: representative images of Elav>P-*htt* from each time point. Scale bar: 50 *μ*m. **B**. *Left:* Schematic depicting site of imaging on a third-instar larva. Intravital images (24, 48, 72 Hours after Hatching) and dissected preparations (96 and 120 Hours after Hatching) of segmental nerve branches (SNB). First row: representative images from Elav>NP-*htt* at each time point. Second row: representative images from Elav>P-*htt* at each time point. Scale bar: 25 *μ*m. **C.** Average RFP Fluorescence intensity (AU) from the VNC for larvae expressing Elav>NP-*htt* vs Elav>P-*htt* across all five timepoints imaged. **D.** Average number of RFP aggregates per 100μm of SNB for larvae expressing Elav>NP-*htt* vs Elav>NP-*htt* across all five timepoints imaged. **E-H:** Data from 120 HAH larvae. **E.** Average RFP aggregate size (μm) of all aggregates of 120 HAH larvae in Elav>NP-*htt* (green) vs Elav>P-*htt* (red). Inset: Elav > NP-htt only. **F.** Average RFP Fluorescence intensity (AU) of all aggregates of 120 HAH larvae between Elav>NP-*htt* (green) vs Elav>P-*htt* (red). Inset: Elav > NP-htt only. **G.** Average aggregate size of 120 HAH larvae in Elav>NP-*htt* vs Elav>P-*htt*. **H.** Average aggregate fluorescence intensity (AU) of 120 HAH larvae between Elav>NP-*htt* and Elav>P-*htt.* * indicates P>0.05, ** P>0.01, *** P>0.001, **** P>0.0001 from a One-way ANOVA.

To track the progression of P-htt aggregation in neurons more thoroughly, the main segmental nerve branches (SNB) projecting from the VNC were imaged (Fig 2 B). These axons contain motor neurons projecting to body-wall muscles ^43^. No detectable htt-aggregates were observed in the SNBs of Elav > NP-htt larvae until 120 HAH (Fig 2 Bi-v, D, One-way ANOVA, F=104.2, P<0.0001). For Elav > P-htt larvae, no htt-aggregation was observed 24 HAH; however, a prolific increase in the number of aggregates, quantified per 100 *μ*m of SNB, was observed 48 HAH (Fig 2 Bvii, D). Subsequently, the 72 HAH timepoint did not show a significant increase compared to 48 HAH, but the 96 and 120 HAH timepoints revealed a significant increase compared to 48 HAH, reaching a plateau in aggregate number per 100 *μ*m after 96 HAH (Fig 2 Bviii-x, D). Statistical differences were observed between the Elav > NP-htt and Elav > P-htt larvae at the 48, 72, 96, and 120 HAH timepoints (Fig 2 D, One-way ANOVA). To examine quantifiable differences between Elav > NP-htt and Elav > P-htt expressing in larvae, a comparison of the two genotypes was conducted at the 120 HAH timepoint. Every SNB aggregate was plotted as a function of its size to demonstrate the profound differences in both number and size (Fig 2 E). Next, each aggregate was plotted as a function of its fluorescence intensity (Fig 2 F). Significant differences were observed between Elav > NP-htt and Elav > P-htt larvae at 120 HAH in the average aggregate size (Fig 2 G, NP: 5.6 ± 3.1, P: 11.2 ± 3.2, T-test, t=2.8, P=0.02) and average fluorescence intensity (Fig 2 H, NP: 401.7 ± 36.2, P: 2248.2 ± 984.7, T-test, t=4.2, P=0.003). Collectively, these results demonstrate a prolific increase in htt-aggregates from the pathogenic construct beginning as early as 24 HAH in the VNC, and 48 HAH in the axons, while comparatively few htt-aggregates are present in the VNC or motor-axons in the non-pathogenic construct.

### Trafficking of organelles in axons of P-and NP-htt expressing third instar larvae

To assess the impact of htt-aggregates on motor neuron (MN) morphology and physiology, we first explored the trafficking of organelles within these SNB using *in vivo* live imaging. Given these MNs are glutamatergic, an assessment of the impact of aggregation on trafficking of synaptic vesicles (SVs) containing the neurotransmitter (NT) glutamate was conducted by expressing a genetically-encoded GFP-tagged version of synaptotagmin 1 (UAS-Syt1-GFP), the well-characterized calcium-sensor for SV-fusion (Fig 3 A) ^45–47^. SV flux (number/min) within the SNB was calculated for Elav > UAS-Syt1-GFP (10.7 ± 3.3, Fig 3 C). Expressing this construct in the NP-htt background did not significantly alter SV flux (10.5 ± 3.8, Fig 3 C, One-way ANOVA, F=27.5, P>0.05). However, when examining the impact of SV flux in flies also expressing P-htt, SV trafficking was significantly impaired (4.7 ± 2.0, Fig 3 C, One-way ANOVA, F=27.5, P<0.0001). To directly examine the impact of htt-aggregation on trafficking, SV flux was calculated up-and downstream of SNB with aggregates exceeding 5 *μ*m^2^ (Fig 3 B). Upstream of an htt-aggregate, SV trafficking was significantly impaired compared to both Elav-Gal4 > UAS-Syt1-GFP, and Elav-Gal4 > UAS-Syt1-GFP, NP-htt (Fig 3 B, C). A kymograph depicts the uncoordinated, back and forth movement of GFP-positive puncta (SVs) upstream (*a*-anterior, *p*-posterior) of a htt-aggregate, and a dramatic reduction in SV movement downstream (Fig 3 Biii). Impact of htt-aggregation in P-htt animals on SV trafficking was most apparent downstream, and significantly different from upstream activation as well as from SV trafficking in NP-htt animals (1.4 ± .9, Fig 3 C, One-way ANOVA, F=27.5, P<0.0001). To localize htt-aggregation more precisely within individual MNs, we attempted to examine the impacts of trafficking within single axons by using MN-Ib-Gal4 and MN-Is-Gal4 drivers (Fig 3 D) ^38^. However, neither driver was strong enough to visualize both htt-aggregates and SV trafficking simultaneously. The number of htt-aggregates in individual axons within SNB, observable with the single MN-driver, MN-Ib-Gal4>P was comparable to what was observed using Elav (Data not shown).

**Fig 3.**
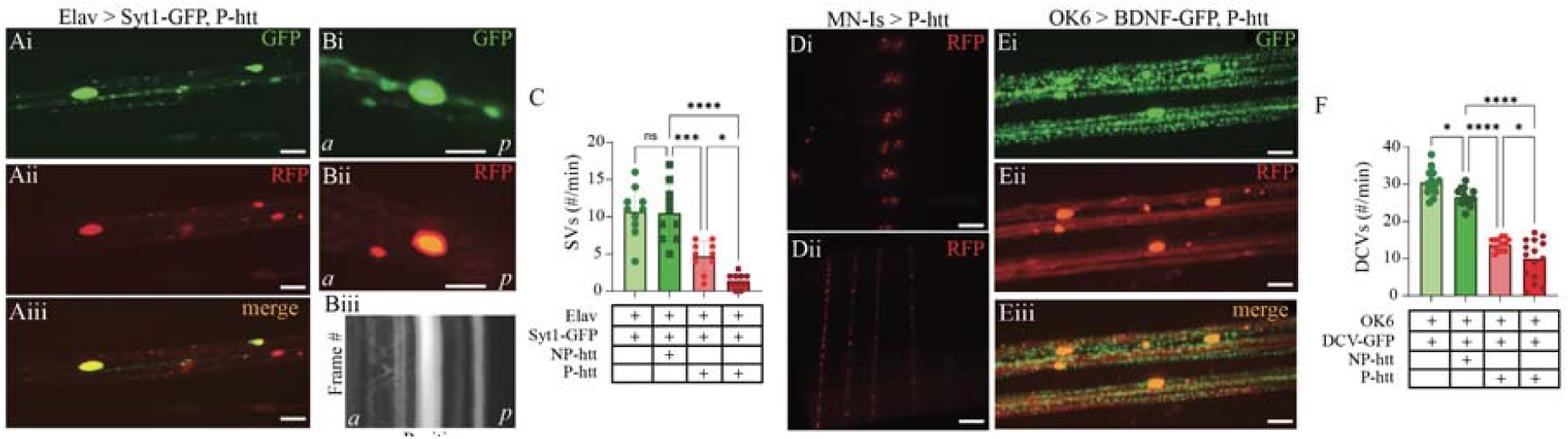
Htt aggregates cause significant trafficking deficits of synaptic vesicles and dense core vesicles. **Ai-iii.** Confocal microscopy images of Elav > Syt1-GFP, P-htt from a segmental nerve branch. Scale bar: 10 µm. **Bi-ii.** Confocal microscopy image of individual htt-aggregate from Elav > Syt1-GFP, htt-P. Scale bar: 5 µm. **Biii.** Kymograph of Syt1-GFP SVs from Bi, highlighting differences in mobility up-and down-stream of aggregate. *a:* anterior/upstream*, p:* posterior/downstream. **C.** Quantification of SV flux (#/min). Genotypes are indicated below figure, + indicates presence of genetic element. Light red: quantification upstream of aggregate, Dark red bar: quantification downstream of aggregate. **Di-Dii.** Fluorescence microscopy image of MN-Is > P-htt in the **(Di**) VNC and (**Dii**) SNBs. Scale bar: 50 µm (Di), 25 µm (Dii). **Ei-iii.** Confocal microscopy images of OK6 > BDNF-GFP, P-htt from a segmental nerve branch. Scale bar: 10 µm. **F.** Quantification of dense core vesicle flux (#/min). Genotypes are indicated below figure, + indicates presence of genetic element. Light red: quantification upstream of aggregate, Dark red bar: quantification downstream of aggregate. * indicates P>0.05, ** P>0.01, *** P>0.001, **** P>0.0001 from a One-way ANOVA.

### Impacts of htt on Dense Core Vesicle trafficking

To further examine the impacts of htt-aggregates on axonal trafficking, we examined the impact on dense core vesicle (DCV) trafficking. DCVs are considerably larger (2-3x) than SV in diameter, and have a dark and granulated core on electron micrographs resulting from dense packing of their cargo, which can contain numerous cell-signaling and regulatory components, including neuropeptides, peptide hormones, growth factors, signaling proteins, biogenic amines and active zone components ^48–53^. Substantial evidence suggests that transport deficits of BDNF are key contributors to HD ^54,55^. Consequently, we created a novel DCV overexpression line tagged with GFP using traditional cloning techniques incorporating the UAS-system to overexpress the neuropeptide brain-derived neurotropic factor (UAS-BDNF-GFP, Fig 3 E). The molecular weight of the construct plus GFP was validated using a Western blot. Kymographs were created to calculate the trafficking metrics of BDNF-tagged DCVs in SNBs, including velocity, flux, and duty cycle. NS expression of BDNF-GFP using Elav-Gal4 revealed a baseline flux calculation of 30.5 ± 3.6 (Fig 3 F). Trafficking of DCVs was significantly impaired in Elav-Gal4 > UAS-BDNF-GFP, NP-htt compared to Elav-Gal4 > UAS-BDNF-GFP (Fig 3 F, One-way ANOVA, F= 101.1, P>0.05). However, DCV trafficking was significant reduced when P-htt was expressed (Fig 3 F, One-way ANOVA, F= 101.1, P<0.001). As observed for SV trafficking, DCV trafficking was impaired to a greater extent downstream of htt-aggregates compared to upstream (Fig 3 F). These results demonstrate that axonal trafficking is significantly reduced only in the presence of P-htt aggregates, and not NP-htt aggregates. Moreover, the effects of htt-aggregates on organelle trafficking are most dramatically reduced downstream of htt-aggregates.

### Htt expression at the neuromuscular junction

Following the observation that organelle trafficking was significantly impaired in P-htt axons, we next sought to examine the downstream or developmental effects of htt-aggregations. The *Drosophila* larval glutamatergic neuromuscular junction (NMJ) has served as an invaluable model glutamatergic synapse for over half a century ^43^. To establish if htt-aggregation was visible at the NMJ, whole-mount immunohistochemical stains for RFP and HRP were conducted. Htt-aggregates were not detected at the NMJ of Elav > NP-htt third instar larvae (Fig 4 Ai). In the 8 immunostains conducted for Elav > P-htt third instar larvae, aggregates were observed at the NMJ innervating muscle fiber (MF) 4 at a single NMJ in MF 4 (Fig 4 Aii). Of note, htt-aggregates were often observed at branching points upstream of the NMJ (data not shown). The total length of MN innervation along the surface of MF 4, a large, flat muscle ideal for NMJ morphology assessments, was measured (Fig 4 A). No significant difference was observed between Elav > NP-htt and Elav > P-htt expressing lines (Fig 4 B, One-way ANOVA, F= 41.1, P>0.05). Two morphologically and physiologically distinct MN-subtypes, MN-Ib (tonic-like) and MN-Is (phasic-like), are located at the larval NMJ. A separate assessment of each MN-subtype was conducted to isolate MN-specific effects of htt-aggregates on innervation length, but no significant differences were observed (Fig 4 B). Next, an assessment of the total bouton number, along with a separation of bouton number per MN-subtype was conducted and no significant differences were observed (Fig 4 C). Lastly, an assessment of the number of active zones (AZs) per bouton was also calculated using an immunostain against a member of the ELKS-family of scaffolding proteins -bruchpilot (brp, Fig 4 Aiii) -and no significant difference between Elav > NP-htt and Elav > P-htt expressing lines was observed (data not shown). Taken together, Elav > NP-htt nor Elav > P-htt expressing lines had any observable effects on NMJ presynaptic structural morphology.

**Fig 4.**
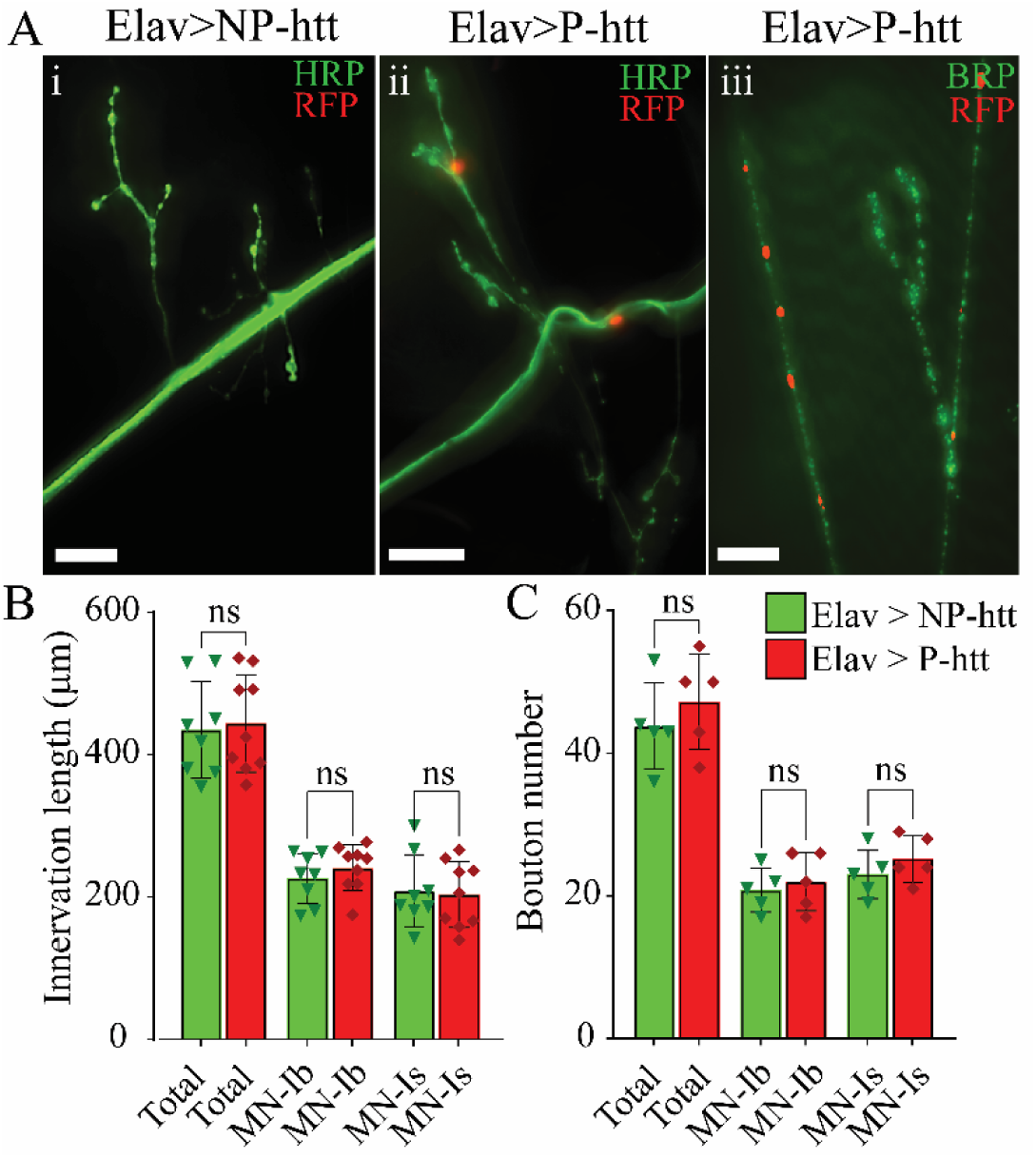
Htt-aggregation does not impact larval NMJ morphology. **A.** Immunohistochemical stain from **i.** Elav-Gal4 > NP-htt and **ii:** Elav-Gal4 > P-htt using horseradish peroxidase (HRP) and red fluorescent protein (RFP). **Aiii.** Immunostain of Elav-Gal4 > UAS-htt-Q138 for bruchpilot (brp) and RFP. Scale bar: 50 *μ*m. **B.** The total innervation length of motor neurons (MNs) along the surface of muscle fiber 4 was quantified for both Elav > P-*htt* and Elav > NP-*htt*. The MNs were further divided into the MN-Ib and MN-Is subtypes. **C.** Total bouton number along the survive of MF4 was quantified for both Elav > P-*htt* and Elav > NP-*htt*. The bouton numbers were further divided assessed by associating them to either MN-Ib or MN-Is subtypes.

### Effects of htt-aggregation on neuromuscular transduction

Although no morphological deficits were observed at the NMJ, electrophysiological recordings were conducted to examine the physiological impacts of NP-and P-htt expression at the NMJ (Fig 5). Unstimulated, sharp-intracellular recordings were initially performed to examine changes in miniature excitatory junctional potential (minis) amplitude and frequency (Fig 5 A). No significant change in mini amplitude was observed between control (Elav), Elav > NP-htt, and Elav > P-htt lines (Fig 5 C, One-way ANOVA, F=2.8, P>0.05). There was also no significant change in mini frequency between the control, Elav > NP-htt, and Elav > P-htt lines (Fig 5 D, One-way ANOVA, F=0.7, P>0.05). Next, low frequency stimulation was applied to severed SNBs via a suction electrode at 0.2 Hz to generate excitatory junctional potentials (EJPs, Fig 5 B). Similar to the effects on minis, no significant deficits in the evoked release of synaptic vesicles were observed between control, Elav > NP-htt, and Elav > P-htt larvae EJPs (Fig 5 E, One-way ANOVA, F=0.7, P>0.05). Collectively, our electrophysiological recordings do not demonstrate any physiological impacts of htt-aggregation on neuromuscular transduction.

**Fig 5.**
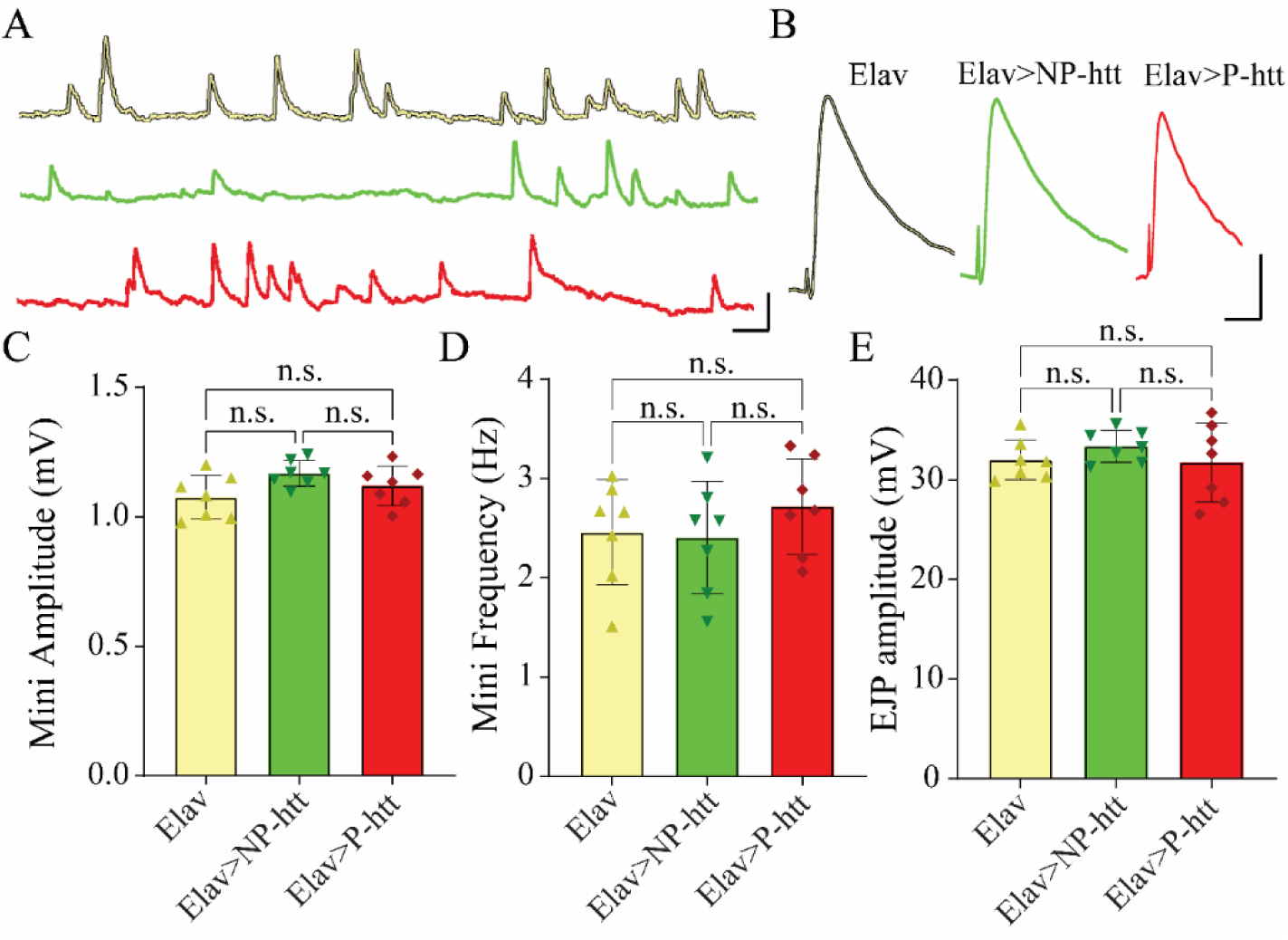
Htt aggregation in motor neurons does not significantly impact NMJ transmission. **A.** Representative trace of Miniature excitatory post junction potentials (MINI) from Elav (yellow), Elav>NP-htt (Yellow), and Elav>P-htt (red). Scale bars: 1 mV, 5 ms. **B.** Representative trace of excitatory post junction potentials (EJP) from Elav (yellow), Elav>NP-htt (Yellow), and Elav>P-htt (red). Scale bars: 15mV, 20 ms. **C.** MINI amplitude (mV) of Elav (Elav), Elav>NP-htt (NP), and Elav>P-htt (P). **D.** MINI frequency (Hz) for of Elav (Elav), Elav>NP-htt (NP), and Elav>P-htt (P). **E.** EJP amplitude (mV) of Elav (Elav), Elav>NP-htt (NP), and Elav>P-htt (P).

### Effects of htt-aggregation on muscle morphology

Next, an assessment of muscle morphology was conducted to examine changes in postsynaptic structure (Fig 6). Immunohistochemical stains to examine changes in gross muscle structure were conducted using the actin-stain phalloidin (Fig 6 A-D). Gross morphological assessments of muscle fiber (MF) 4 from third instar larvae Elav, Elav > NP-htt, and Elav > P-htt were conducted and no significant differences were observed for MF length (Fig 6 Dii, One-way ANOVA, F=0.5, P=0.6), MF width (Fig 6 Diii, One-way ANOVA, F=4.1, P>0.05), and MF area (Fig 6 Div, One-way ANOVA, F=1.6, P=0.2). To examine possible changes in postsynaptic ultrastructure, a fluorescence profile line feature was used within Nikon Elements to measure fluorescence peaks along a discrete line of a minimum length of 200 µm as a precise estimation of muscle sarcomere and I-band lengths (Fig 6 Ei.) ^40^. Immunohistochemical stains from Elav > NP-htt and Elav > P-htt larvae did not show any significant changes in either sarcomere length (Fig Eiii, T-Test, t=0.46, P=0.65) or I-band length (Fig Eiv, T-test, t=0.82, P=0.43). Lastly, an immunostain using the nuclear stain DAPI was conducted to examine changes in muscle nuclei structure and morphology (Fig 6 F). Stains from Elav > NP-htt and Elav > P-htt expressing larvae did not show any significant differences in nuclei number (Fig 6 Fii, T-test, t=1.21, P=0.26), nuclei area (Fig. 6 Fiii, T-test, t=0.84, P=0.40), and internuclear distance (Fig 6. Fiv, T-test, t=1.55, P=0.13). Collectively there does not appear to be any postsynaptic effects of Elav > NP-htt or Elav > P-htt aggregation at the NMJ.

**Fig 6.**
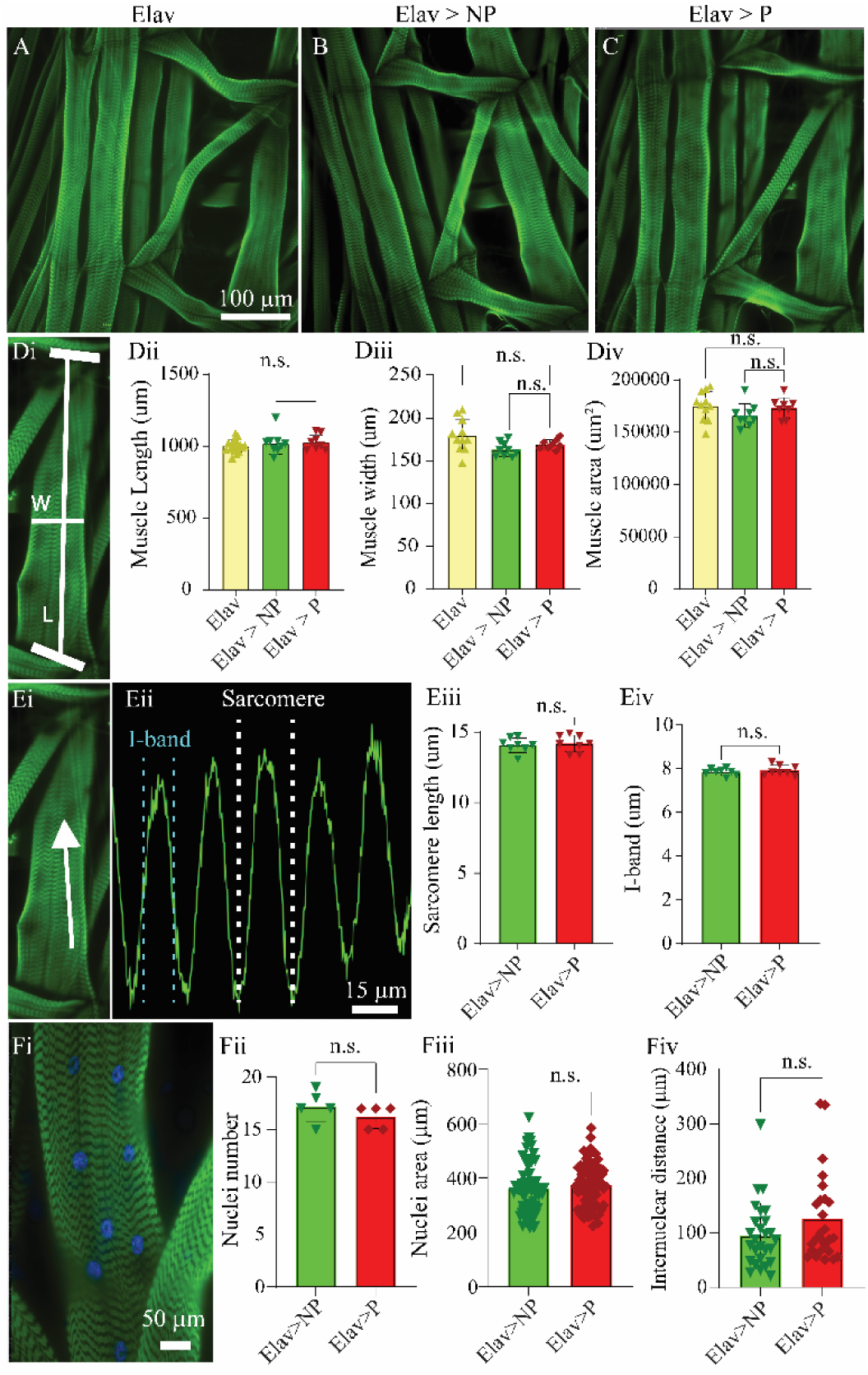
Htt-aggregation does not impact muscle ultrastructure. Immunostain using phalloidin (actin) of an abdominal hemisegment highlighting longitudinal muscles 4, 5, 6, 7, 12, and 13 from **A.** Elav, **B.** Elav > NP-htt, and **C.** Elav > P-htt. Scale bar: 100 *μ*m. **Di.** Immunostain of MF12 demonstrating how muscle width (W) and length (L) measurements were obtained. Quantification of gross muscle length (**Dii**), width (**Diii**), and area (**Div**) for Elav, Elav > NP-htt, and Elav > P-htt. **Ei.** Immunostain of MF12 showing how muscle ultrastructure was determined using a fluorescence line plot. Along white arrow indicates the location and direction of ultrastructural analysis. **Eii.** Fluorescence intensity profile line for GFP (actin). Dashed white and teal lines indicate how sarcomere and I-band measurements were calculated respectively. Scale bar: 15 *μ*m. Quantification for muscle (**Eiii)** I-band length and (**Eiv**) sarcomere length measurements from Elav > NP-htt, and Elav > P-htt. **Fi.** Immunostain for MF12 with phalloidin and DAPI. Scale bar: 50 *μ*m. Quantification of nuclei number (**Fii**), nuclei area (**Fiii**), and internuclear distance (**Fiv**) from Elav > NP-htt, and Elav > P-htt.

### Htt-aggregation effects on excitation-contraction coupling machinery

The MNs contained within the SNBs projecting from the VNC transmit descending patterned outputs, driving muscle contraction in larval body-wall muscles responsible for larval peristalsis ^42,53,56^. To examine the impact of NP-and P-htt expression in the NS on mechanisms of the excitation-contraction coupling machinery, a force transducer system was used to measure changes in muscle force production from semi-intact larvae (Fig 7 A). With the brain removed, the motor neurons were activated via electrical stimulation using a series of frequencies within the range commonly output from the VNC to generate a force-frequency curve for each genotype ranging from 1-150 Hz (Fig 7 B) ^42^. Elav > NP-htt third instar larvae did not show any differences in muscle contraction compared to Canton S. controls, however, Elav > P-htt larvae had significantly reduced muscle force production (Fig 7 C). The amplitude of contractions generated at 40 Hz were reduced by 34.6 ± 5.1 % in Elav > P-htt compared to Elav > NP-htt larvae (Fig 7 C, *peak force*). Interestingly, the rise and decay time constant (Fig 7 C, *τ*_rise_ and *τ*_decay_) were also slower for Elav > P-htt compared to Elav > NP-htt or Canton S. controls (*τ*_rise_: 33 ± 13% increase, and *τ*_decay_: 88 ± 31% increase). Force frequency curves generated in Figure 7 Di (300 ms) show profound changes in muscle performance in Elav > P-htt expressing larvae compared to Elav > NP-htt and controls. Elav > P-htt larvae did not show any observable muscle contractions until the stimulation frequency exceeded 10 Hz, while observable contractions were elicited in Elav > NP-htt and Canton S. larvae at 1 Hz (Fig 7 Di). A 30-39% decrease in muscle force production was observed between 10-40 Hz, the aspect of the force-frequency curve with greatest plasticity ^42^. Contraction force was significantly reduced at all stimulation frequencies below 100 Hz that were examined between the two constructs (Fig 7 Di). A well-documented leftward shift in the force-frequency curve is observed by increasing the duration of the intraburst stimulation ^42^. Figures 7 Dii-iv, show force-frequency curves for larvae with 600, 750, and 900 ms duration simulation. By increasing the duration of the intraburst frequency, greater temporal summation occurs, resulting in greater larval force generation, and a steady decline in the differences in force production between Elav >NP-htt and Elav > P-htt expressing larvae. Despite this leftward shift, significant differences exist between Elav > P-htt and Elav > NP-htt from 10-40 Hz for all simulation durations (Fig 7 D).

**Fig 7.**
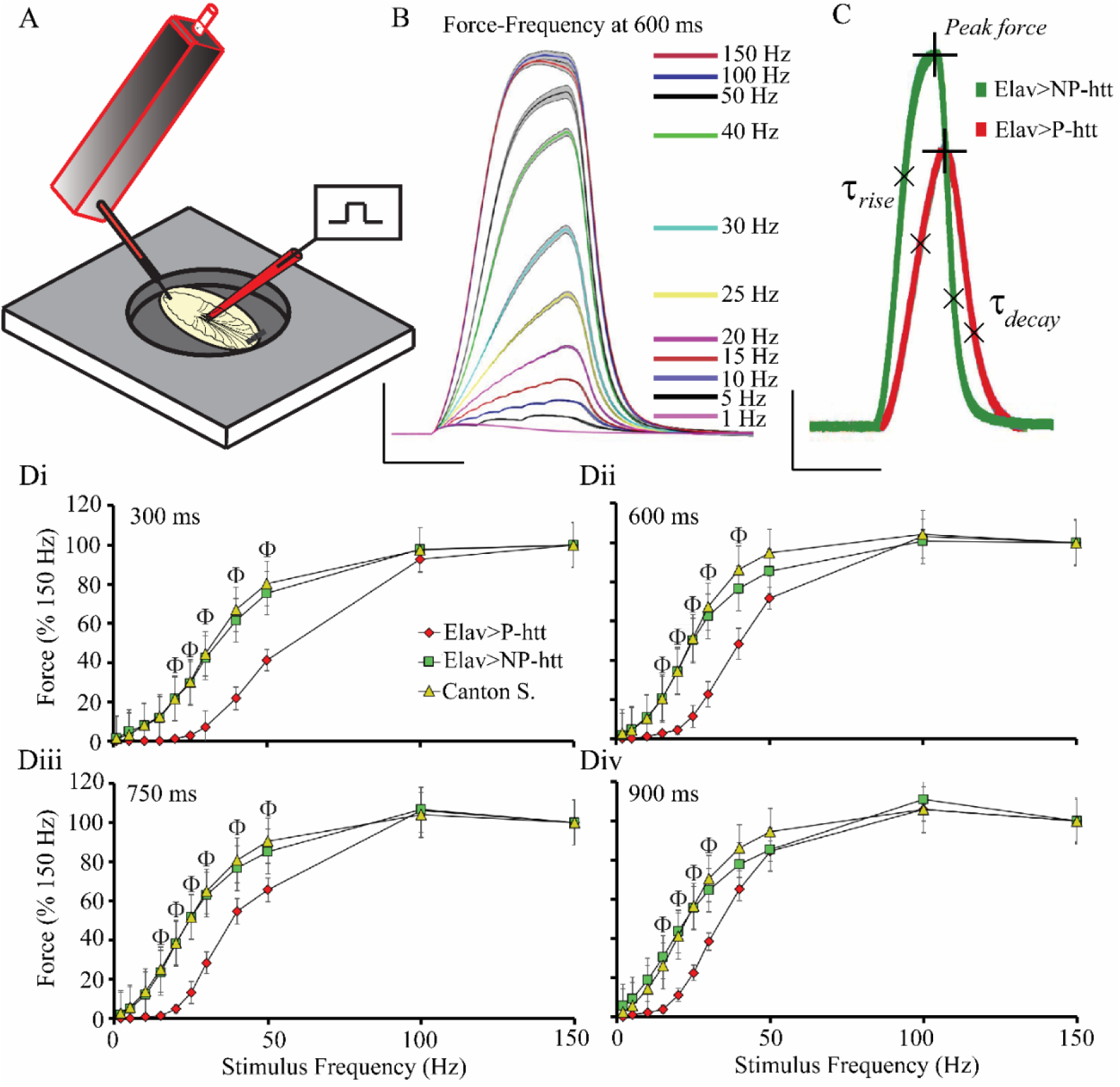
P-htt expression in motor neurons significantly reduces neuromuscular transduction. **A.** Schematic of larval force transducer setup. **B.** Representative force-frequency recordings from third-instar larvae. Six replicate contractions from each of the 11 stimulation frequencies were averaged and plotted with a 95% confidence interval. **C.** Representative traces from Canton S. control and P-htt larvae with peak force, rise-and decay-tau (*τ*) shown. Scale bars: 1mN, 750 ms. **D.** Force-frequency plots for 4 different stimulation durations 300 ms (Di), 600 ms (Dii), 750 ms (Diii), 900 ms (Div). Φ: indicates significant differences between Elav>NP-htt and Elav > P-htt.

### Larval crawling and human-htt expression

Given the profound effects of P-htt expression on muscle contraction, an assessment of larval crawling was conducted to determine if alterations in muscle contraction biology manifested as changes in locomotor or crawling behavior. In sets of 10 larvae at a time, 100 larvae from each genotype were placed inside a dark box devoid of light, illuminated and recorded using infrared light. Their crawling behavior was tracked using CTRAX software, and assessed for velocity, and distance travelled (Fig 8 A) ^57^. UAS-NP-htt, UAS-P-htt, Elav, and Elav > NP-htt larvae all displayed similar crawling behavior (Fig 8 B, C, One-way ANOVA, P>0.05). Larva expressing Elav > P-htt showed a significant reduction in crawling velocity compared the other 4 genotypes (Fig 8 B, One-way ANOVA, F=36.63, P<0.0001). Elav > P-htt expressing larvae also showed a significant reduction in the distance travelled over a period of 30 seconds of continuous crawling (Fig 8 C, One-way ANOVA, F=28.01, P<0.0001).

**Fig 8.**
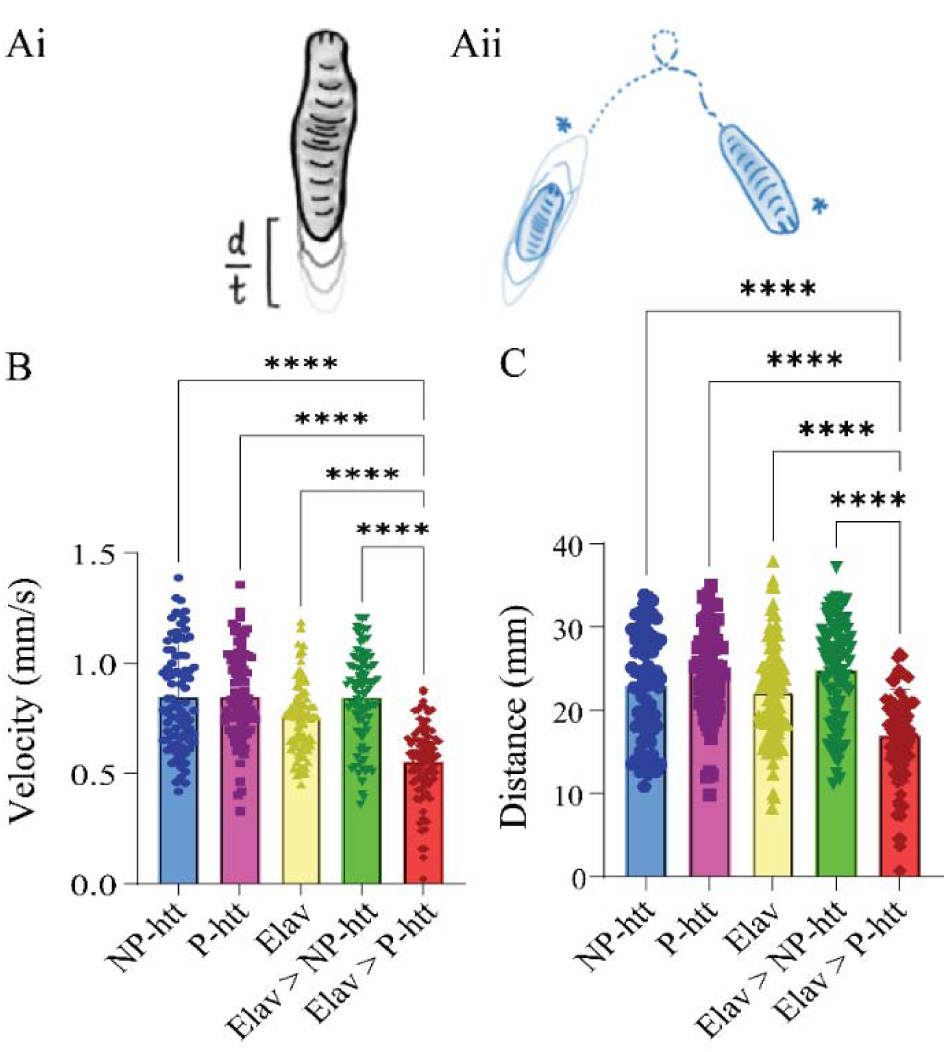
Pathogenic htt-expression in the NS significantly impacts larval crawling. **A.** Schematic depicting different crawling patterns quantified: velocity (i) and distance (ii). **B.** Velocity (mm/s) is significantly decreased in Elav > P-htt compared to Elav > NP-htt and all controls. **C.** Distance traveled (mm) over 30 seconds is significantly decreased in Elav > P-htt compared to Elav > NP-htt and all controls. * indicates P>0.05, ** P>0.01, *** P>0.001, **** P>0.0001 from a One-way ANOVA.

### Htt expression in the wings of adult Drosophila reveal similar patterns to the larval NS

A limitation in using larvae as a model for investigations of neurodegeneration is the relatively short lifespan. To complement our current approach, we sought to develop a model to investigate cellular htt-aggregation across the lifespan of adult *Drosophila*. Expressing P-htt using Elav has previously been demonstrated to result in a significantly reduced adult lifespan ^33^. Furthermore, imaging of htt-aggregation in the adult CNS is technically challenging. A screen for Gal4-drivers in the wing was conducted to circumvent both these obstacles, and appl-Gal4 was identified to express in the L1 vein of the wing (Fig 9 A,)^34^. Adults expressing appl > NP-htt and appl > P-htt were isolated on the day of eclosion, and the right wing of unique animals were imaged transcutaneously every 5 days for 50 days. Small and infrequent aggregates were observed in the wings of appl > NP-htt adults while similarly sized, yet more numerous aggregates were observed in the wings of 5-day old appl > P-htt animals (Fig 9 Bi, iv, C, D). For appl > NP-htt animals, no significant increase was observed for subsequent 5-day intervals from 5-50 days, indicating no significant increase the aggregate number (Fig 9 C, One-way ANOVA, F=14.44, P=0.0001). For appl > P-htt animals, a significant difference was observed between day 5 and days 45 -50, demonstrating a significant increase with age (Fig 9 C). Most notably, a significant difference was observed in the number of htt aggregates between appl > NP-htt and appl > P-htt at each time-point, except day 10, showing significantly more htt-aggregates in the animals expressing the pathogenic construct (Fig 9 C). For each aggregate, both the area and fluorescence intensity were also quantified (Fig 9 D-E). No significant increase in aggregate area was observed for appl > NP-htt animals over the 50 days investigated (Fig D, One-way ANOVA, F=1795, P<0.0001). Interestingly, the size of aggregates in appl > P-htt wings did not show an increase between days 5 and 10, however, a steady and gradual significant increase in the size of aggregates was observed in subsequent time points (Fig 9 D). Significant differences in aggregate area were observed between the two genotypes after day 20, revealing significantly larger aggregates in the pathogenic version of htt (Fig 9 D). The fluorescence intensity of appl > NP-htt wing aggregates did not show any significant increases, with one exception at day 20, where a large spike in fluorescence intensity was observed (Fig 9 E, One-Way ANOVA, F=190.6, P<0.0001). Contrary to aggregate size, wing aggregates from appl > P-htt animals showed a steady increase in intensity from days 5-to-20, following by a dramatic decrease (Fig 9 E). Significant differences in fluorescence intensity were observed between the two genotypes for the first 3 time-points (Fig 9 E).

**Fig 9.**
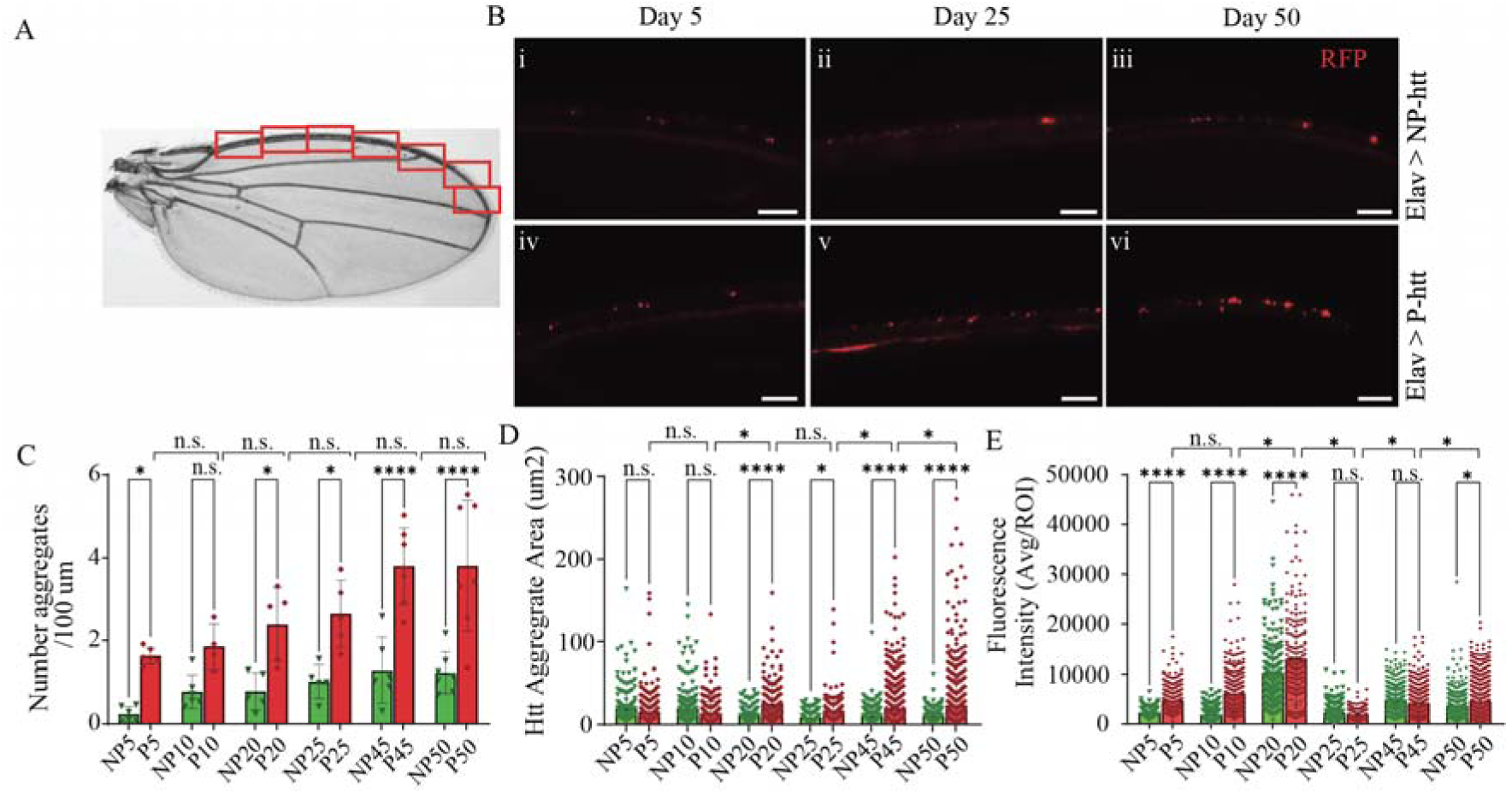
Modelling htt-aggregation in the adult wing reveals significant increases in the number and size of aggregates from flies expressing pathogenic htt. **A.** Bright-field microscope image of an adult *Drosophila* wing, highlighting the location of images along the L1 vein in red boxes. Scale bar: 50 *μ*m. **B.** Fluorescence microscopy images from the wings of Elav > NP-htt (Bi-iii) and Elav > P-htt (Biv-vi) from 3 time points, day 5, 25, and 50. Scale bar: 50 µm. Quantification of the (**C)** number of aggregates per 100 µm, (**D**) aggregate area, and (**E**) fluorescence intensity from Elav > NP-htt and Elav > P-htt. * indicates P>0.05, ** P>0.01, *** P>0.001, **** P>0.0001 from a One-way ANOVA.

### Expression of htt in adults shows significant reduction in lifespan

To assess the impacts of NP-and P-htt expression in the CNS of adult flies, initially a lifespan assay was conducted, again using Elav-Gal4. For the 3 control lines, the average lifespan of the 200 flies assessed for each genotype was over 80 days (Fig 10 A, NP-htt, P-htt, Elav-Gal4). Flies expressing Elav > NP-htt did not show any reduction in lifespan compared to controls; 50% lethality was 58 days in Elav > NP-htt flies compared to 66.0, 78.1, and 64.7 days in Elav, P-htt and NP-htt flies respectively. 90% lethality at 75.3 days in Elav > NP-htt flies compared to 81.2, 98.4, and 95.5 for Elav, P-htt and NP-htt flies respectively (Fig 10 A). However, Elav > P-htt expressing flies showed a drastically reduced lifespan compared to the other four genotypes, with 50% lethality at 23.7 days, and 90% lethality at 28.6 days (Fig 10 A). These results are consistent with previous reports that expression of pathogenic human htt constructs in the NS of adult flies significantly reduces lifespan ^33^.

**Fig 10.**
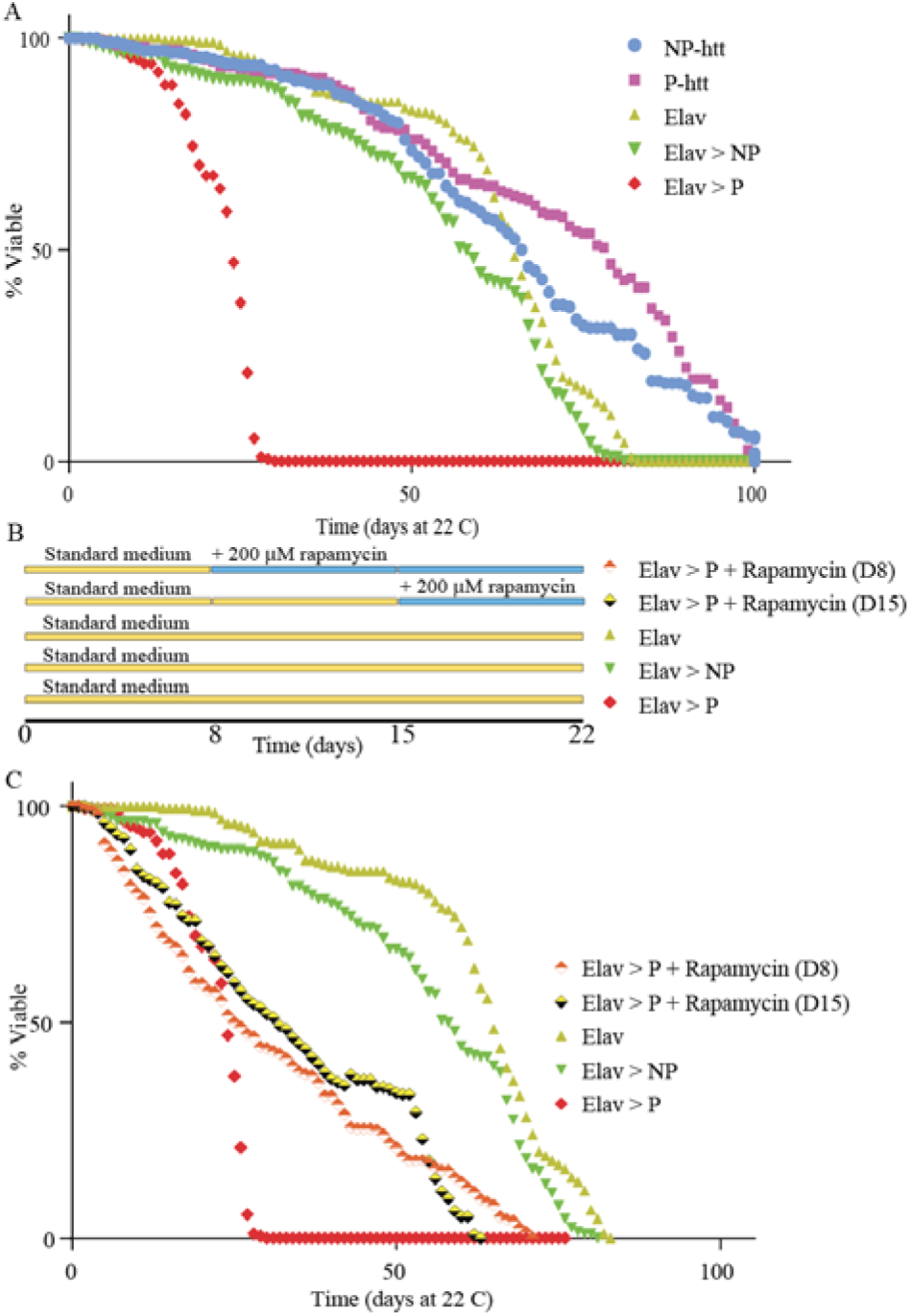
Expression of htt aggregates significantly reduces adult lifespan, which is partially rescued with rapamycin feeding. **A.** Lifespan (% viability) was tracked for adults reared at 22°C for 5 different genotypes plotted as a function of time in days. **B.** Schematic showing the feeding schedule of rapamycin for the experiments in C. **C.** Lifespan (% viability) was tracked for adults reared in 22°C for 3 different genotypes and Elav > P-htt fed standard medium supplemented with 200 µM rapamycin 8 days (D8) and 15 days (D15) post eclosion.

### Effects of htt expression on lifespan can be rescued by oral administration of the mTOR inhibitor rapamycin

Targeting the mTOR pathway has been shown to rescue lifespan deficits linked to polyQ disorders ^13–15^. Herein, we fed adult flies the mTOR inhibitor, Rapamycin, to upregulate the mTOR dependent autophagy pathway, beginning either 8 or 15 days post-eclosion (Fig 10 B, ref). Administration of rapamycin (200 µM rapamycin in standard food) 8-and 15-days post eclosion to adult Elav > P-htt flies dramatically rescued their lifespan: 50% lethality at 26 and 31.6 days, and 90% viability at 62.4 and 57.5 days, beginning feeding at 8 and 15-days post-eclosion respectively (Fig 10 C). Supplementing our standard media with 200uM Rapamycin was able to more than double that lifespan of Elav > P-htt flies.

## Discussion

Modeling Huntington’s Disease in *Drosophila* facilitates a robust exploration of the molecular, cellular, physiological, and behavioral impacts of the disease. Herein, we utilized the extensive molecular and genetic toolkits in *Drosophila* to track *in vivo* progression of htt-aggregates throughout development and assessed the downstream effects of expanded polyQ-htt aggregation within the central and peripheral nervous systems. *In vivo* imaging facilitated tracking of htt-aggregate size, number, and fluorescence intensity as a function of time across precise windows of development. Animals with pathogenic htt exhibited aggregates of significantly greater number, size, and intensity at nearly all developmental time points. These aggregates caused severe trafficking impairments for organelles, up-and downstream of aggregates. While no obvious downstream effects were observed on the morphology of pre-and postsynaptic structures in the NMJ, or electrophysiological recordings, a distinct reduction in muscle force production was observed despite normal muscle ultrastructure. In our model of HD, animals expressing pathogenic htt exhibit locomotor deficits in both crawling velocity and distance. To longitudinally study HD in adult *Drosophila*, we developed a novel model for htt-aggregation in adult wings, which demonstrated similar patterns of htt-aggregate development to larvae. Lastly, pathogenic expression in the NS of flies results in significantly reduced viability, though supplementation of rapamycin, an mTOR pathway inhibitor, partially rescued fly viability and extended their lifespan.

A neuropathological hallmark of human HD is intracellular aggregation of N-terminal htt fragments, suggesting HD neuropathology is mediated by aberrant proteolytic cleavage or removal of htt fragments ^58^. While numerous proteases have been demonstrated to cleave N-terminal htt, proteolysis at the caspase-6 site is an important pathogenic event in HD ^59,60^. *In vitro* expansion of N-terminal htt fragments enhances cytotoxicity but site-directed mutagenesis of the caspase-6 cleavage site of N-terminal htt reduces cytotoxicity, suggesting inhibition of caspase cleavage of htt may be neuroprotective ^61,62^. Caspase-6 mediated N-terminal fragments of human htt were found to form aggregates throughout the soma, axons, and occasionally synapses of motor neurons ^44^. In mouse models of HD, htt fragments formed aggregates progressively over time in the CNS, mirroring observations made in neuropathological studies of asymptomatic striatal, cerebral, and cerebellar brain slices from human HD patients ^17,18,20,24,60^. In our study using a *Drosophila* model of HD, noticeable aggregates began forming in the somas in less than 24 HAH and continued to accumulate significantly until plateauing at 72 HAH. Similar effects were observed in the axons, increasing steadily from 48-96 HAH, then plateauing. The dynamics of htt-aggregate formation have been studied, both *in vitro* and *in vivo* models of HD, revealing similar patterns of mobility, proliferation, and accumulation ^33,63^. *In vitro* studies expressing fluorescently tagged exon one mhtt-94Q fragments in mouse embryonic stem cells have identified fast diffusion, dynamic clustering, and stable aggregation as three distinct dynamic states ^63^. Similarly, *Drosophila* studies expressing fluorescently tagged exon one htt-138Q fragments identified rapid diffusion of fragments in axons, but faster incorporation of mobile htt-Q138 following aggregate formation and hasty reformation following photobleaching ^33^. Our results also support these observations showing that initially fragmented P-htt accumulates into numerous, small aggregates, which increase in size exponentially over a few days, eventually reaching a plateau. Additionally, we demonstrated that SV and DCVs colocalized with htt-aggregates, suggesting that not only do htt-aggregates quickly accumulate new htt protein, but also act as an indiscriminate protein sink on a seconds-to-minutes timescale. Taken together, preventing, limiting, and clearing htt fragment aggregation could be a critical aspect of HD neuropathology and a target for treatment strategies.

The endogenous biological role of human htt remains unclear but previous studies have implicated htt in numerous processes including: as a scaffold protein, transcription regulation, neurogenesis, macroautophagy, cargo transport, and synaptic assembly and plasticity ^64–70^. These numerous roles are unsurprising, given the large number of protein-protein interaction domains within htt as well as its documented interactions with hundreds of proteins ^71,72^. Furthermore, the subcellular localization of htt is fluid and dynamic, potentially due to an intrinsic capacity for intracellular compartment-dependent conformational changes ^126^. Antibodies recognizing different epitopes within htt reveal unique subcellular labelling including the nucleus, endosomes, ER, Golgi, axons, and synapses ^73–79^. Electron microscopy of htt reveals strong interactions between SVs and microtubules ^80^. Here we demonstrate profound impacts of htt-aggregates on axonal trafficking of organelles. Both neurotransmitter-containing SVs and neuropeptide-containing DCVs are trafficked along the axonal microtubules via canonical molecular motor machinery ^81,82^ While a direct interaction between dynein/dynactin has been identified, htt is known to be a critical scaffold for both molecular motors: kinesin and the dynein/dynactin complex ^68,76,83^. Huntington-associated-protein-1 (HAP1), one of the earliest identified binding partners of htt, directly binds to Kinesin-light-chain (KLC) and Dynactin p150^glued^, as evidenced by mutations or alterations in HAP1 expression are known to significantly impair axonal trafficking dynamics Pathogenic htt has been demonstrated to exhibit a higher binding affinity for HAP1, leading to breakdowns in the intracellular trafficking of SVs, DCVs, and mitochondria ^83–85^. Furthermore, a critical stretch of 17 amino-acids at the start of N-terminal of htt regulates mitochondrial and ER localization ^86^. Htt is also known to facilitate clathrin-mediated vesicle trafficking into endosomes via its interaction with Huntingtin-interacting protein (HIP1) ^87^. In dendrites, htt has been found directly interacting with dynein to mediate trafficking of TrkB-containing vesicles, representing a more extensive transport role for htt ^88^. We investigated organelle trafficking both up-and downstream of aggregates to distinguish the endogenous role of htt in trafficking organelles from the effect of aggregates on downstream trafficking. Trafficking of SVs and DCVs was significantly impaired upstream of htt-aggregates, supporting the critical role of htt in regulating molecular trafficking. Interestingly, an even greater impact of P-htt was observed on trafficking downstream of aggregates, suggesting the effects of aggregation outweigh the defects in its role in supporting molecular transport proteins. Aggregation not only impairs trafficking by acting as a sink for newly trafficking htt, but it severely limits downstream trafficking of organelles. Collectively, these results suggest expansion of the polyQ region in exon one of htt alters the endogenous functionality of the protein, its association with other critical motor proteins, and leads to aggregation that severely impairs organelle trafficking, and other cellular functions.

Vesicular transport deficits in BDNF are known to play a critical role in HD ^54,89^. Consequently, we explored the impacts of P-htt on the trafficking dynamics of BDNF and found a dramatic reduction in its molecular transport. Studies have shown that decreasing BDNF production in cortical neurons results in degeneration of the striatum, but restoring BDNF secretion promotes neuronal survival ^90–92^. While BDNF is not directly produced in striatal neurons, it is transported to the striatum by corticostriatial-projecting neurons in the cortex ^93^. Although several models of HD pathology implicate pathogenic htt-mediated trafficking deficits in late-onset symptoms, numerous other adult-onset neurodegenerative disease are attributed to transport breakdowns, indicating a strong link between trafficking deficits and neurodegeneration pathology generally ^94^.

Surprisingly, htt aggregation and trafficking deficits observed herein did not alter the morphology or electrophysiological properties of the neuromuscular junction. As such, evidence of neuronal degeneration downstream of numerous, large htt aggregates was not overt. Presynaptic motor neurons formed seemingly wild-type innervation along the surface of body wall muscles. At the ultrastructural level, no changes in bouton number or AZ density were observed in either glutamatergic motor neuron subtypes. Similarly, postsynaptic muscle fibers were structurally normal at both gross and ultrastructural organizational levels. It is important to note the presence of satellite boutons, indicative of a putative compensatory mechanism deployed by the neuron to mitigate transmission deficits, consistent with previous reports ^95^. Indeed, insoluble aggregates formed in the axoplasm do not lead to neurodegeneration, but evidence suggests that nuclear aggregates could contribute to the neurodegenerative processes ^20,23,96^. Nuclear aggregates have been shown to sequester cAMP response element-binding protein (CREB), a critical transcription factor for mediating cell survival, contributing significantly to neurodegeneration observed in HD and other neurological disorders ^97^. Studies have demonstrated that soluble forms of polyQ-htt, rather than insoluble aggregates, contribute to neurodegeneration despite the severe reductions in lifespan observed when large aggregates are expressed in the NS ^33,98^. Indeed, studies in mice show neurodegeneration in the absence of htt-aggregation ^23,99,100^. Furthermore, despite striatal neurons being the site of highest degree of neurodegeneration in human patients, aggregates observed in these neurons are exceedingly rare ^20^. While multiple studies have demonstrated that nuclear translocation of htt increases cytotoxicity and neuropathy, evidence also shows widespread nuclear htt without exhibiting neurotoxicity ^129,130^. It is therefore difficult to determine whether expanded polyQ N-terminal fragments of htt are sufficient for disease progression, or if another fragment of htt is needed for nuclear localization and/or neurodegeneration. Nevertheless, while no obvious effects were observed to support either developmental perturbation or expanded polyQ htt fragment aggregate-induced changes in NMJ morphology, profound effects on muscle contractility were observed.

One of the earliest clinically diagnosed symptoms of HD is chorea, involuntary jerking or writhing movements typically associated with altered CNS functionality ^101^. However, recent work has focused on the effects of HD in peripheral tissues, including muscles ^102,103^ . Using a microforce transducer system, we observed significantly reduced muscle contraction force in animals expressing the expanded polyQ htt fragment. Muscle force production was most severely impaired during 10-50 Hz motor neuron stimulation, previously identified as the most plastic and modifiable stimulation frequency ^42^. Descending motor output recorded from intact larvae show stimulation frequencies ranging from 1-150 Hz, while fictive muscle contraction recordings demonstrated 25-30 Hz as the most common during rhythmic peristalsis ^42^. Shorter duration stimulation bursts (200-300 ms) were more severely impacted, but significant effects were observed during pronounced 900 ms duration stimulations. The rise and decay time kinetics were also severely impacted, indicating potential issues with calcium liberation, or other aspects of the excitation-contraction coupling machinery ^42^. Given these well-defined descending output frequencies underlying muscle contraction and locomotion, it is not surprising low-frequency stimulation EJPs did not reveal a significant change. High frequency stimulation underlying locomotion triggers many well-characterized dynamic processes at the bouton ^104–106^. High frequency dependent cycling of the readily releasable, recycling, and reserve pools and new supplies of synaptic materials from the soma are tightly regulated processes critical for normal muscle ^127^. Furthermore, the presence of aggregates in motor neurons at critical branching points observed herein may cause AP failure during high frequency stimulation ^128^. Thus, the molecular machinery and metabolic demand during fictive contraction recordings is likely a better representation of the downstream impacts of htt-fragment aggregates on neuromuscular transduction.

Given that atrophy and skeletal muscle wasting is a common hallmark of HD, it is not surprising that transcriptional changes in skeletal tissues were comparable to those observed in different brain regions ^103^ . Several mechanisms have been implicated in skeletal muscle HD pathology, including mitochondrial abnormalities, cytotoxic aggregation, and reduced heat shock transcription factor 1 ^107–110^. However, the impacts of HD on neuromuscular transmission and muscle physiology remain poorly investigated ^110^. Importantly, improving muscle function in murine models delays HD onset and improves lifespan ^111^, and exercise in humans has been shown to improve countless metrics of skeletal muscle pathophysiology in HD patients ^112^. Taken together, investigations of HD on skeletal muscles, and targeting skeletal muscle for therapies are important future directions.

Treatments targeting aggregation centered around protein stabilization, administration of small molecule inhibitors, and upregulating cell-clearance mechanisms show promising improvements in aggregate reduction and mitigating functional deficits. ^113–115^ Considerable research has implicated a strong link between HD and altered macroautophagy, highlighted by a severe reduction in neuronal capacity to degrade misfolded proteins ^116^. Given that P-htt misfolds and forms insoluble aggregates, targeting autophagy was of particular interest in our current investigation. P-htt has been shown to alter cellular levels of Argonaute 2, a critical protein of the RISC complex modulating post transcriptional regulation of gene expression, caused by reduced autophagy leading to both neurodegeneration and functional deficits ^27^. Furthermore, P-htt has been shown to sequester autophagosomes, further emphasizing htt-aggregates as sink ^117^. Rapamycin, a macrolide mTOR inhibitor, has been demonstrated to both delay and rescue neuronal dysfunction in models of HD ^118–120^. Rapamycin treatment upregulates Atg1 which is critical in the recruitment of other Atg proteins responsible for autophagosome synthesis ^121,122^. Additionally, inhibition of TORC1 by rapamycin depresses protein synthesis in the cell and leads to activation of ULK1 (Atg1 homolog) ^123^. ULK1, integral to autophagosome formation. Unsurprisingly, administration of rapamycin has been shown to enhance P-htt clearance leading to rescue of locomotor function of *Drosophila* ^120^. Previous studies have shown rapamycin to have antiproliferative properties, associated with an effect on lifespan and viability ^124,125^. We explored the effect of these properties by supplementing the food of adult *Drosophila* expressing P-htt with 200 uM rapamycin, which resulted in an effective doubling of their lifespan. Compiling all the evidence, exploration of htt aggregation on disrupting the RNA silencing and autophagy machineries may provide a key link to delaying or preventing aggregate proliferation on cellular fate.

HD remains a challenging disorder to study due to the complex role of the htt protein in various cellular processes. In this study, we reveal the role of P-htt in disrupting the molecular motor machinery and its tendency to form aggregates that indiscriminately trap axonal cargo. Additionally, we observe significant deficits in muscle contractility and locomotory behavior reminiscent of chorea, despite no noticeable deformities in NMJ formation or muscle ultrastructure. Targeted therapies, whether pharmacological or genetic, will depend on a comprehensive understanding of the pathophysiological effects of P-htt aggregates and their downstream targets, which may extend beyond the nervous system.

The authors declare no competing interests.

## Acknowledgements

Research supported by the National Institute of General Medical Sciences of the National Institutes of Health under Award Number R15GM155985 to KGO. Generous financial support from Middle Tennessee State University to KGO. The authors would like to thank Professor Troy Littleton for generously providing the UAS-NP-htt (mRFP-HttQ15), UAS-P-htt (mRFP-HttQ138), and UAS-Syt1-GFP constructs, as well as the GluRIII antibody. We would also like to thank Dr. Yulia Akbergenova for her insights on the manuscript.

